# A replication-competent vesicular stomatitis virus for studies of SARS-CoV-2 spike-mediated cell entry and its inhibition

**DOI:** 10.1101/2020.05.20.105247

**Authors:** M. Eugenia Dieterle, Denise Haslwanter, Robert H. Bortz, Ariel S. Wirchnianski, Gorka Lasso, Olivia Vergnolle, Shawn A. Abbasi, J. Maximilian Fels, Ethan Laudermilch, Catalina Florez, Amanda Mengotto, Duncan Kimmel, Ryan J. Malonis, George Georgiev, Jose Quiroz, Jason Barnhill, Liise-anne Pirofski, Johanna P. Daily, John M. Dye, Jonathan R. Lai, Andrew S. Herbert, Kartik Chandran, Rohit K. Jangra

**Author notes:** These authors made equivalent contributions (M. Eugenia Dieterle, Denise Haslwanter). Corresponding authors (R.K.J.), (K.C.), (A.S.H.).

## Abstract

There is an urgent need for vaccines and therapeutics to prevent and treat COVID-19. Rapid SARS-CoV-2 countermeasure development is contingent on the availability of robust, scalable, and readily deployable surrogate viral assays to screen antiviral humoral responses, and define correlates of immune protection, and to down-select candidate antivirals. Here, we describe a highly infectious recombinant vesicular stomatitis virus bearing the SARS-CoV-2 spike glycoprotein S as its sole entry glycoprotein that closely resembles the authentic agent in its entry-related properties. We show that the neutralizing activities of a large panel of COVID-19 convalescent sera can be assessed in high-throughput fluorescent reporter assay with rVSV-SARS-CoV-2 S and that neutralization of the rVSV and authentic SARS-CoV-2 by spike-specific antibodies in these antisera is highly correlated. Our findings underscore the utility of rVSV-SARS-CoV-2 S for the development of spike-specific vaccines and therapeutics and for mechanistic studies of viral entry and its inhibition.

## Introduction

A member of the family *Coronaviridae*, severe acute respiratory syndrome coronavirus-2 (SARS-CoV-2) is the causative agent of the ongoing coronavirus disease 2019 (COVID-19) pandemic that emerged in Wuhan City, China in late 2019 (Wu et al., 2020a). With more than 4.8 million confirmed cases and at least 320,000 deaths in over 216 countries/areas/territories as of May 19, 2020, the global scale and impact of COVID-19 is unparalleled in living memory (Dong et al., 2020; WHO Situation Report May 19th, 2020). To date, mitigation strategies have relied largely on physical distancing and other public health measures. Although treatments with some small molecule inhibitors and with convalescent plasma have received approvals for emergency use; and vaccines, antivirals, and monoclonal antibodies are being rapidly developed, no FDA-approved countermeasures are currently available.

The membrane-enveloped virions of SARS-CoV-2 are studded with homotrimers of the spike glycoprotein (S), which mediate viral entry into the host cell (Bosch et al., 2003); (Walls et al., 2020; Wrapp et al., 2020). S trimers are post-translationally cleaved in the secretory pathway by the proprotein convertase furin to yield *N*– and *C*–terminal S1 and S2 subunits, respectively. S1 is organized into an *N–*terminal domain (NTD), a central receptor-binding domain (RBD), and a *C*–terminal domain (CTD). S2 bears the hallmarks of a ‘Class I’ membrane fusion subunit, with an *N*–terminal hydrophobic fusion peptide, *N*– and *C*–terminal heptad repeat sequences, a transmembrane domain, and a cytoplasmic tail (Walls et al., 2020; Wrapp et al., 2020).

The S1 RBD engages the viral receptor, human angiotensin-converting enzyme 2 (hACE2) at the host cell surface (Hoffmann et al., 2020; Wang et al., 2020; Zhou et al., 2020). Receptor binding is proposed to prime further S protein cleavage at the S2’ site by the transmembrane protease serine protease-2 (TMPRSS2) at the cell surface, and/or by host cysteine cathepsin(s) in endosomes. S2’ cleavage activates S2 conformational rearrangements that catalyze the fusion of viral and cellular membranes and escape of the viral genome into the cytoplasm (Hoffmann et al., 2020).

The S glycoprotein is the major antigenic target on the virus for protective antibodies (Rogers et al., 2020; Walker et al., 2020; Wu et al., 2020b), and is thus of high significance for the development of vaccines and therapeutic antibodies. Plasma derived from human convalescents and replete with such antibodies has shown early promise as a COVID-19 treatment, and it is currently being evaluated in clinical trials of antiviral prophylaxis and therapy (Casadevall and Pirofski, 2020). Considerable efforts are also being aimed at the identification and deployment of S glycoprotein-specific neutralizing monoclonal antibodies (mAbs) (Cao et al., 2020; Pinto et al., 2020; Rogers et al., 2020; Walker et al., 2020; Wu et al., 2020b; Zost et al., 2020).

A key requirement for the rapid development of such vaccines and treatments with convalescent plasma, small-molecule inhibitors, and recombinant biologics is the availability of safe, robust, and faithful platforms to study S-glycoprotein inhibition with high assay throughput. Given limited access to biosafety level-3 (BSL-3) containment facilities required to safely handle SARS-CoV-2, researchers have turned to surrogate viral systems that afford studies of cell entry at biosafety level-2 (BSL-2) and facilitate rapid inhibitor screening through the use of fluorescence or luminescence-based reporters. These include retroviruses, lentiviruses, or vesiculoviruses ‘pseudotyped’ with SARS-CoV-2 S and competent for a single round of viral entry and infection (Lei et al., 2020; Nie et al., 2020; Ou et al., 2020; Pu et al., 2020; Tan et al., 2020; Xiong et al., 2020). However, these single-cycle pseudotyped viruses are typically laborious to produce and challenging to scale up, yield poorly infectious preparations, and suffer background issues in some cases due to contamination with viral particles bearing the orthologous entry glycoprotein (e.g., low levels of vesicular stomatitis virus (VSV) pseudotypes bearing VSV G).

In contrast to the single-cycle pseudotypes, replication-competent recombinant VSVs (rVSVs) encoding the heterologous virus entry glycoprotein gene(s) *in cis* as their only entry protein(s) are easier to produce at high yields and also afford forward-genetic studies of viral entry. We and others have generated and used such rVSVs to safely and effectively study entry by lethal viruses that require high biocontainment (Caì et al., 2019; Jae et al., 2013; Jangra et al., 2018; Kleinfelter et al., 2015; Maier et al., 2016; Raaben et al., 2017; Whelan et al., 1995; Wong et al., 2010). Although rVSVs bearing the S glycoprotein from SARS-CoV(Fukushi et al., 2006a, 2006b; Kapadia et al., 2005, 2008) and the Middle East respiratory syndrome coronavirus (MERS-CoV) (Liu et al., 2018) have been developed, no such systems have been described to date for SARS-CoV-2.

Here, we generate a rVSV encoding SARS-CoV-2 S and identify key passage-acquired mutations in the S glycoprotein that facilitate robust rVSV replication. We show that the entry-related properties of rVSV-SARS-CoV-2 S resemble those of the authentic agent and use a large panel of COVID-19 convalescent sera to demonstrate that the neutralization of the rVSV and authentic SARS-CoV-2 by spike-specific antibodies is highly correlated. Our findings underscore the utility of rVSV-SARS-CoV-2 S for the development of spike-specific vaccines and antivirals and for mechanistic studies of viral entry and its inhibition.

## Results

### Identification of S gene mutations that facilitate robust rVSV-SARS-CoV-2 S replication

To generate a replication-competent rVSV expressing SARS-CoV-2 S, we replaced the open-reading frame of the native VSV entry glycoprotein gene, *G*, with that of the SARS-CoV-2 *S* (Wuhan-Hu-1 isolate) (Fig. 1A). We also introduced a sequence encoding the enhanced green fluorescent protein (eGFP) as an independent transcriptional unit at the first position of the VSV genome. Plasmid-based rescue of rVSV-SARS-CoV-2 S generated a slowly replicating virus bearing the wild-type S sequence. Five serial passages yielded viral populations that displayed enhanced spread. This was associated with a dramatic increase in the formation of syncytia (Fig. 1B and Fig. S1) driven by S-mediated membrane fusion (data not shown). Sequencing of this viral population identified nonsense mutations that introduced stop codons in the *S* glycoprotein gene (amino acid position C1250* and C1253*), causing 24- and 21-amino acid deletions in the S cytoplasmic tail, respectively. SΔ24 and SΔ21 were maintained in the viral populations upon further passage, and SΔ21 in all plaque-purified isolates, highlighting their likely importance as adaptations for viral growth. Viral population sequencing after four more passages identified two additional mutations, L517S and P812R in S1 and S2, respectively, whose emergence coincided with more rapid viral spread and the appearance of non-syncytium-forming infectious centers (Fig. 1B, passage 5). Pelleted viral particles from clarified infected-cell supernatants incorporated the S glycoprotein, as determined by an S-specific ELISA (Fig 1C).

**Fig 1.**
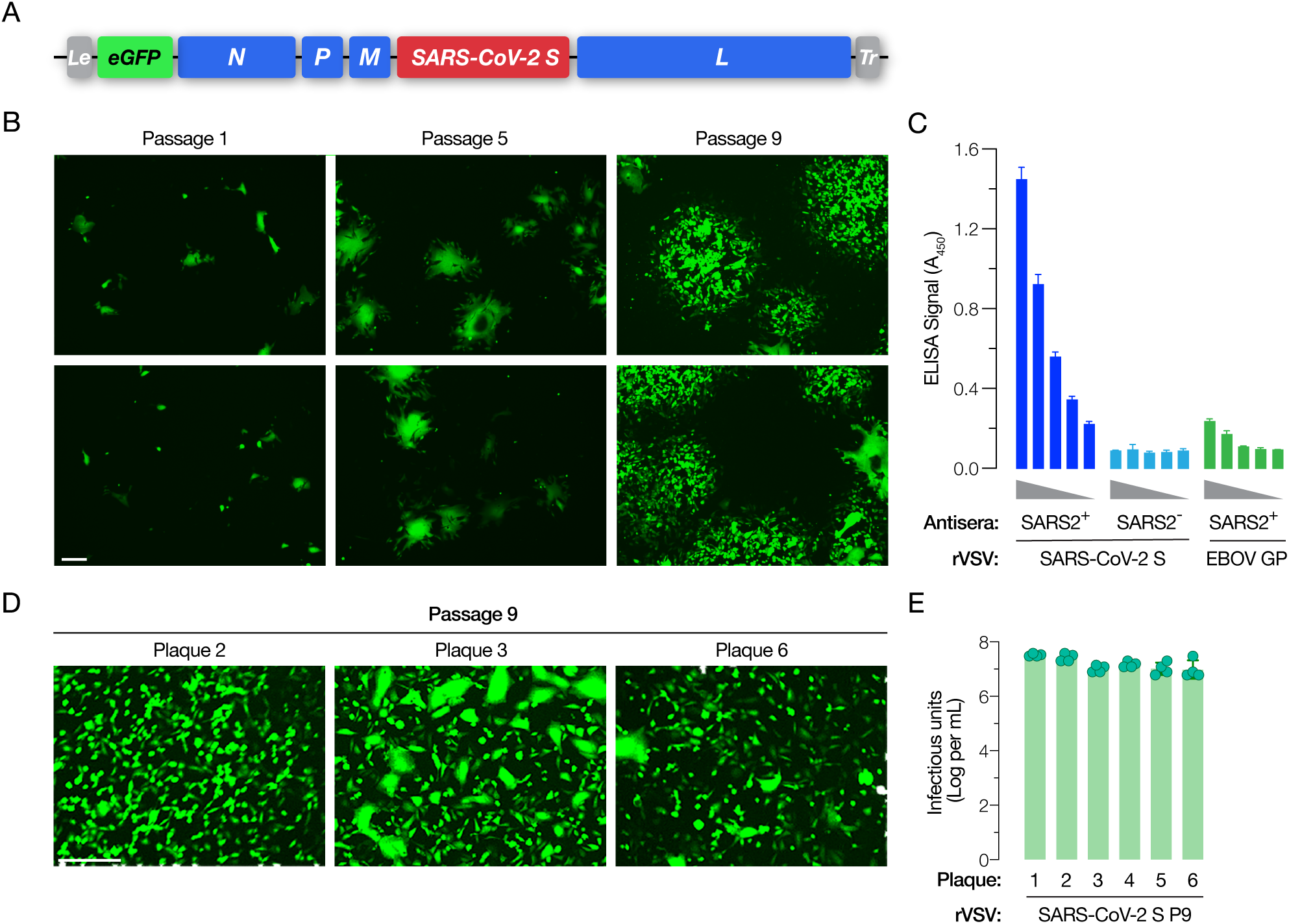
Generation of a recombinant vesicular stomatitis virus (rVSV) bearing the SARS-CoV-2 spike (S) glycoprotein. **(A)** Schematic representation of the VSV genome, in which its native glycoprotein gene has been replaced by that encoding the SARS-CoV-2 S protein. The VSV genome has been further modified to encode an enhanced green fluorescent protein (eGFP) reporter to easily score for infection. **(B)** Infectious center formation assay on Vero cells at 24 h post-infection showing growth of the rVSV-SARS-CoV-2 S after the indicated number of rounds of serial passage of the passage #1 virus (carrying wild-type (WT) S sequences) on Huh7.5.1 cell line (scale bar = 100 µm). Two representative images for each virus passage, showing infected cells in pseudo-colored in green, from one of the two independent experiments are shown here. **(C)** Incorporation of SARS-CoV-2 S into rVSV particles captured on an ELISA plate was detected using antiserum from a COVID-19 convalescent donor (average ± SD, n = 12 from 3-4 independent experiments). Serum from a COVID-19-negative donor and rVSVs bearing Ebola virus glycoprotein (EBOV GP) were used as negative controls (average ± SD, n = 6 from 2 independent experiments). **(D)** Representative images showing Vero cells infected with plaque #2, #3 and #6 viruses at 16 h post-infection (scale bar = 100 µm). **(E)** Production of infectious virions at 48 h post-infection from Vero cells infected with the indicated plaque-purified viruses. Titers were measured on Vero cells overexpressing TMPRSS2 (n = 4, from two independent titrations).

We next sequenced six plaque-purified viral isolates derived from the passage 9 (P9) population. All of these viral clones bore the SΔ21 deletion in the S cytoplasmic tail and spread without much syncytia formation (Fig. 1D). Interestingly, all of these isolates contained three amino acid changes at S-glycoprotein positions other than 517 or 812—W64R, G261R, and A372T—in addition to the *C*–terminal SΔ21 deletion (Table S1). Five of the six isolates also contained mutations H655Y or R685G. Importantly, peak titers of all these viral isolates ranged between 1–3×10^7^ infectious units per mL (Fig. 1E), suggesting that the mutations they share (or a subset of these mutations) drive rVSV-SARS-CoV-2 S adaptation for efficient spread in tissue culture with little or no syncytium formation.

**Fig S1.**
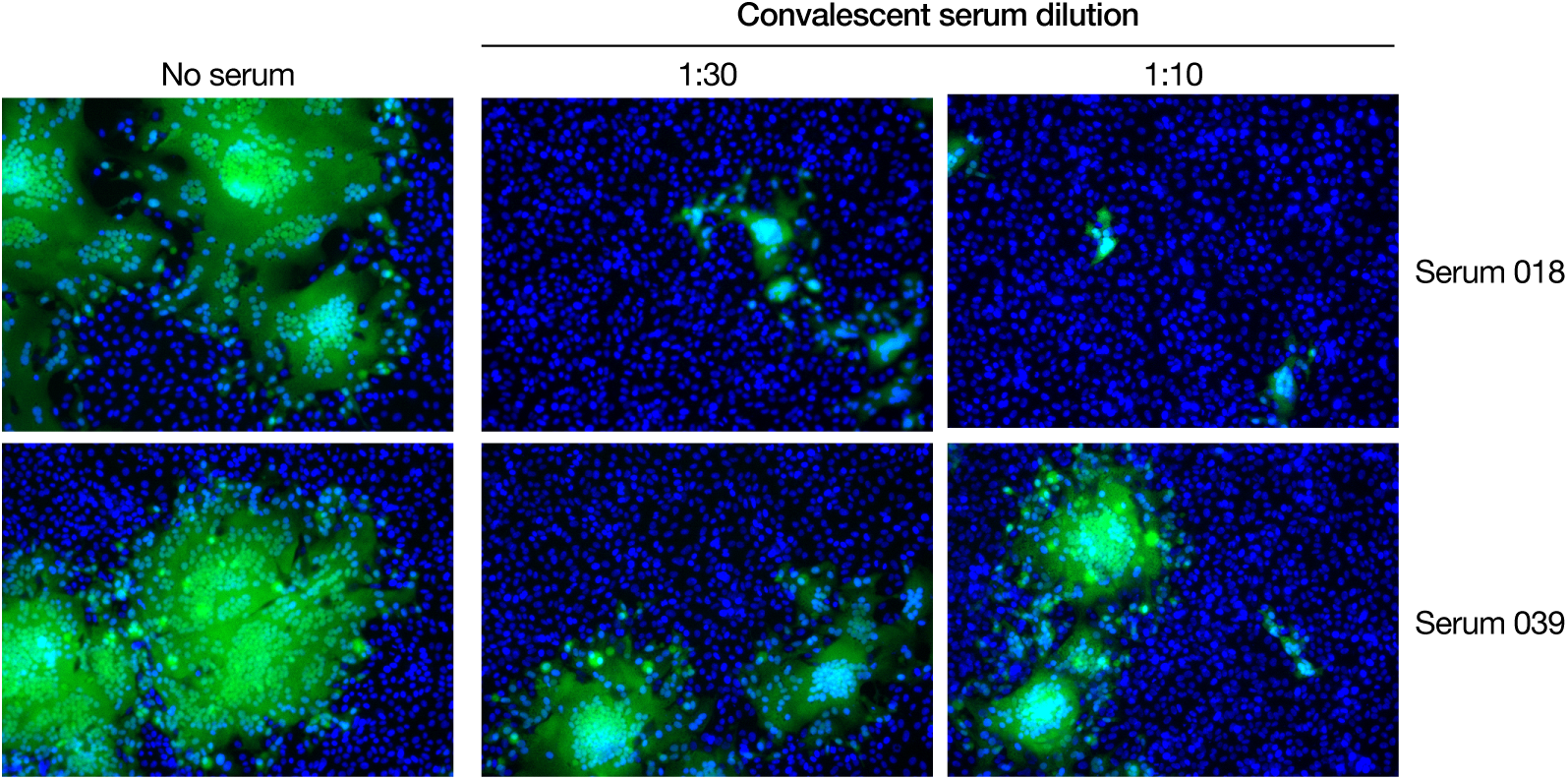
Inhibition of syncytium formation by sera from convalescent donors. Vero cells were infected with pre-titrated amounts of rVSV-SARS-CoV-2. At 2 h post-infection, cells were washed with PBS to remove residual virus and then exposed to the indicated dilutions of each convalescent serum. Cells were fixed, their nuclei counterstained, and they were imaged for syncytium formation by eGFP expression at 16 h post-infection. Representative images from one of two independent experiments are shown.

**Table S1:**
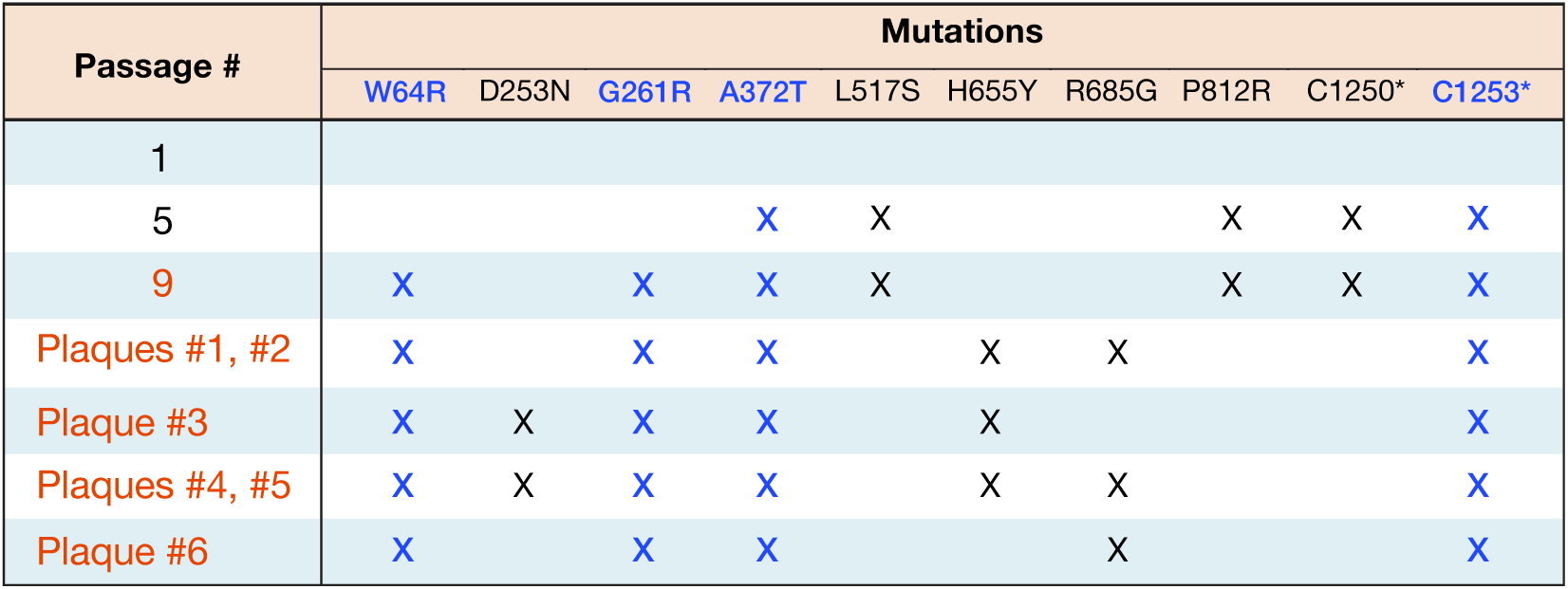
Spike missense and nonsense mutations acquired by rVSV-SARS-CoV-2 S during serial passage

| Passage # | Mutations |  |  |  |  |  |  |  |  |  |
| --- | --- | --- | --- | --- | --- | --- | --- | --- | --- | --- |
|  | W64R | D253N | G261R | A372T | L517S | H655Y | R685G | P812R | C1250* | C1253* |
| 1 |  |  |  |  |  |  |  |  |  |  |
| 5 |  |  |  | X | X |  |  | X | X | X |
| 9 | X |  | X | X | X |  |  | X | X | X |
| Plaques #1, #2 | X |  | X | X |  | X | X |  |  | X |
| Plaque #3 | X | X | X | X |  | X |  |  |  | X |
| Plaques #4, #5 | X | X | X | X |  | X | X |  |  | X |
| Plaque #6 | X |  | X | X |  |  | X |  |  | X |

### rVSV-SARS-CoV-2 S entry is cysteine cathepsin-dependent

SARS-CoV-2 entry in cells has been shown to be dependent on the proteolytic activity of acid-dependent endosomal cysteine cathepsins, including cathepsin L ((Hoffmann et al., 2020; Wang et al., 2020)). Accordingly, we tested the effects of chemical inhibitors of cysteine cathepsins on rVSV-SARS-CoV-2 S infection. Pretreatment of cells with NH_4_Cl, an inhibitor of endosomal acidification, reduced entry by rVSVs bearing SARS-CoV-2 S or the Ebola virus glycoprotein (EBOV GP) in a dose-dependent manner (Fig. 2A). However, S-mediated entry was comparatively less sensitive to NH_4_Cl than that by EBOV GP (Fig. 2A). Next, we tested cysteine cathepsin inhibitors E-64 (Fig. 2B) and FYdmk (Fig 2C). Pre-treatment of cells with both of these compounds also inhibited S-mediated entry, albeit with reduced sensitivity relative to that observed for EBOV GP-dependent entry (Fig. 2B-C). Together, these findings confirm that rVSV-SARS-CoV-2 S resembles the authentic agent in its requirements for endosomal acid pH and cysteine cathepsins. They also suggest a reduced dependence on these host factors for entry by SARS-CoV-2 S relative to EBOV GP, a model glycoprotein known to fuse in late endo/lysosomal compartments following extensive endosomal proteolytic processing.

**Fig 2.**
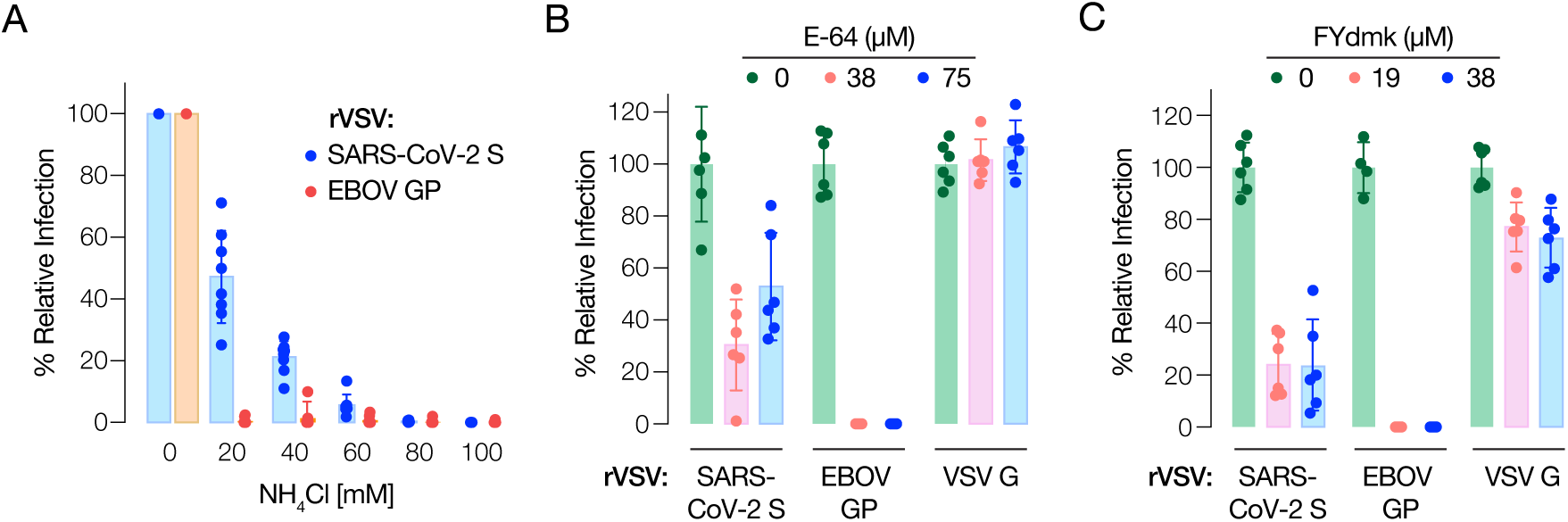
rVSV-SARS-CoV-2 S infection requires the activity of cysteine cathepsin proteases. **(A)** Huh7.5.1 cells pre-treated for 1 h at 37°C with the indicated concentrations of NH4Cl were infected with pre-titrated amounts of rVSVs bearing SARS-CoV-2 S or EBOV GP. Infection was scored by eGFP expression at 16–18 h post-infection (average ± SD, n = 8 from 2 independent experiments). **(B)** Vero cells pre-treated for 90 min at 37°C with the indicated concentrations of pan-cysteine cathepsin inhibitor E-64 were infected with pre-titrated amounts of rVSVs bearing SARS-CoV-2 S, EBOV GP, or VSV G and scored for infection as above (average ± SD, n = 6 from 3 independent experiments, except n = 4 from 2 independent experiments for EBOV GP). **(C)** Vero cells pre-treated for 90 min at 37°C with the indicated concentrations of cathepsin L/B inhibitor FYdmk were infected with pre-titrated amounts of rVSVs bearing SARS-CoV-2 S, EBOV GP or VSV G. Infection was scored as above (average ± SD, n = 6 from 3 independent experiments).

### Human ACE2 is required for rVSV-SARS-CoV-2 S entry

SARS-CoV-2 uses hACE2 as its entry receptor (Letko et al., 2020; Shang et al., 2020b, 2020a). Baby hamster kidney (BHK21) cells do not express detectable levels of ACE2 protein and are resistant to SARS-CoV-2 entry (Hoffmann et al., 2020; Wang et al., 2020). Concordantly, we observed no detectable infection by rVSV-SARS-CoV-2 S in BHK-21 cells (Fig. 3A, top left). By contrast, BHK-21 cells transduced to express hACE2 (Fig. 3A, top right) were highly susceptible to rVSV-SARS-CoV-2 S (Fig. 3A, bottom right and Fig. 3B).

**Fig 3.**
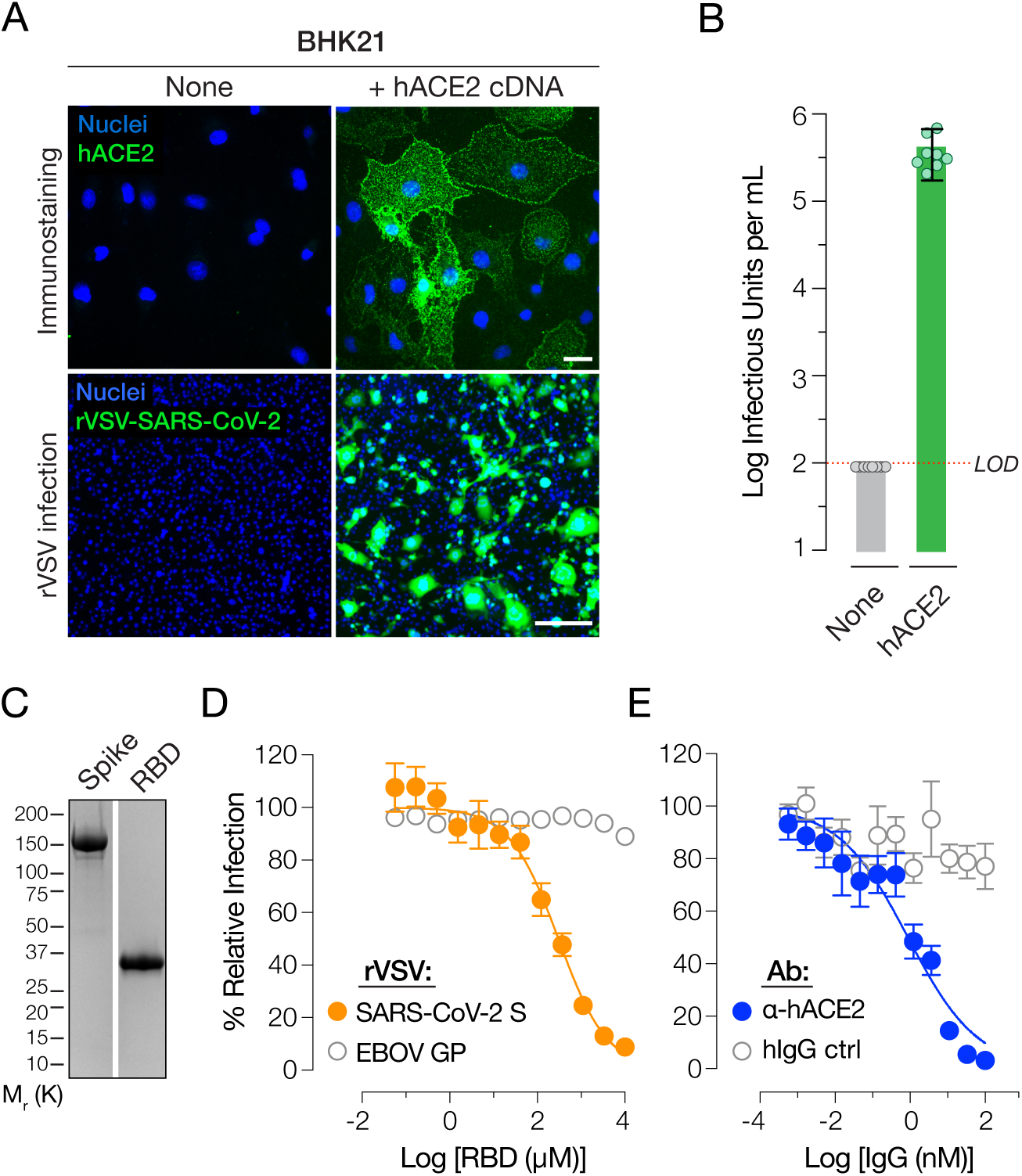
rVSV-SARS-CoV-2 S infection requires human ACE2. **(A)** Naïve (None) baby hamster kidney (BHK21) cells or cells transduced with a retrovirus carrying human ACE2 cDNA (+ hACE2 cDNA) were immunostained for hACE2 expression using an anti-ACE2 antibody. Cells were imaged by fluorescence microscopy. The hACE2 signal is pseudo-colored green (top panel, scale bar = 20 µm). These cells were also exposed to serial 5-fold dilutions of rVSV-SARS-CoV-2 S and infection was scored by eGFP expression (bottom panel, scale bar = 50 µm). Representative images from one of 3 independent experiments are shown. **(B)** Enumeration of eGFP-positive green cells (Average ± SD, n = 8 from 3 independent experiments). LOD: limit of detection. **(C)** Recombinant, Ni-NTA–affinity purified S1-S2 ectodomain (Spike) or the receptor binding domain (RBD) of the SARS-CoV-2 S protein were subjected to SDS-PAGE and Coomassie staining. A representative image from one of two independent purification trials is shown here. **(D)** Monolayers of Huh7.5.1 cells were pre-incubated with serial 3-fold dilutions of the purified RBD for 1 h at 37°C and then infected with pre-titrated amounts of rVSVs bearing SARS-CoV-2 S or EBOV GP. At 16–18 h post-infection, cells were fixed, nuclei counter-stained with Hoechst-33342, and infection (eGFP expression) was scored by fluorescence microscopy. It is represented as % relative infection [no RBD = 100%, Average ± SEM, n = 8 from 3-4 (rVSV-SARS-CoV-2 S) or n = 4 from 2 (rVSV-EBOV GP) independent experiments]. **(E)** Monolayers of Huh7.5.1 cells pre-incubated for 1 h at 37°C with 3-fold serial dilutions of anti-human ACE2 antibody or negative control (hIgG) were infected with pre-titrated amounts of rVSV-SARS-CoV-2 S. Infection was scored as above and is represented as % relative infection [no antibody = 100%, Average ± SD, n = 8 from 3-4 independent experiments].

To directly establish an entry-relevant interaction between rVSV-SARS-CoV-2 S and hACE2, we expressed and purified the spike RBD (Fig. 3C) and pre-incubated it with target cells. RBD pre-treatment inhibited rVSV-SARS-CoV-2 S entry in a specific and dose-dependent manner (Fig. 3D). Moreover, pre-incubation of cells with an hACE2-specific mAb, but not an isotype-matched control mAb, potently abolished rVSV-SARS-CoV-2 S entry (Fig. 3E). These findings provide evidence that rVSV-SARS-CoV-2 S entry and infection, like that of the authentic agent, require spike RBD-hACE2 engagement.

### S protein-targeting antibodies in COVID-19 convalescent sera specifically account for rVSV-SARS-CoV-2 S neutralization

Prior to examining the performance of rVSV-SARS-CoV-2 S in neutralization assays with human antisera, we sought to establish a specific role for interaction between anti-spike antibodies in these sera and the VSV-borne S protein. Accordingly, we first evaluated the reactivity of two sera with rVSV-neutralizing activity (Fig. S2) against viral particles by ELISA. Both sera specifically recognized rVSV-SARS-CoV-2 S particles (Fig. 4A) and were also shown to be reactive against a purified, trimeric preparation of the spike glycoprotein (Wrapp et al., 2020, Bortz et al., manuscript in preparation). Further, serial pre-incubation of each serum with purified S immobilized on a high-binding plate depleted its capacity to inhibit rVSV-SARS-CoV-2 S infection to a degree that was commensurate with the amount of S-specific antibodies it contained (Fig. 4B). By contrast, parallel pre-incubations with blocked plates had little or no effect (Fig. 4B). These results indicate that S glycoprotein-targeting antibodies in COVID-19 convalescent sera specifically mediate rVSV-SARS-CoV-2 S neutralization.

**Fig 4.**
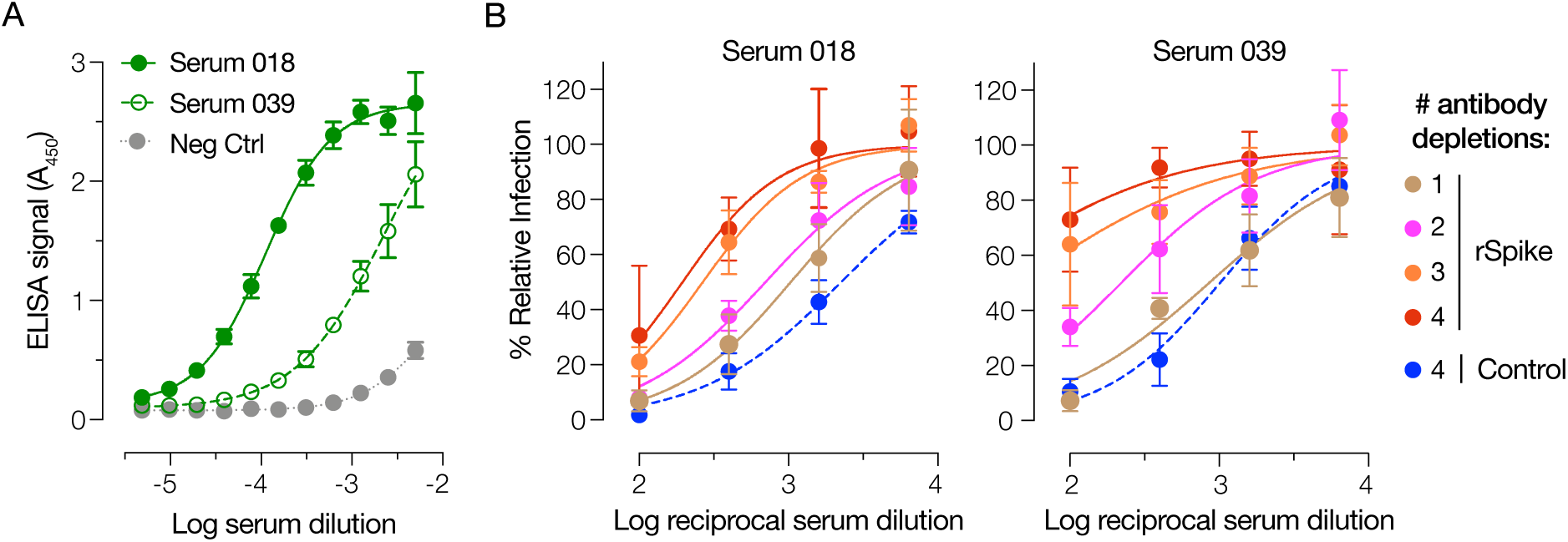
rVSV-SARS-CoV-2 S neutralization is mediated by S glycoprotein-targeting antibodies in human antisera. **(A)** ELISA plates coated with rVSV-SARS-CoV-2 S were incubated with serial 2-fold dilutions of serum 18, serum 39, or negative control serum. Bound S-specific antibodies were detected with an anti-human HRP-conjugated secondary antibody (average ± SD, n = 4 from 2 independent experiments). **(B)** Pre-titrated amounts of serum 18 and serum 39 were sequentially incubated with SARS-CoV-2 S-coated high-binding plates to deplete S-specific antibodies. Capacity of the depleted sera (and control sera incubated with only the blocking agent) to neutralize rVSV-SARS-CoV-2 S was then estimated by incubating pre-titrated amounts of rVSV at the indicated dilutions of sera at 37°C for 1 h prior to infecting monolayers of Huh7.5.1 cells. Cells were scored for infection as above (average ± SD, n = 4 from 2 independent experiments).

### The susceptibilities of rVSV-SARS-CoV-2 S and authentic SARS-CoV-2 to antibody-mediated neutralization are highly correlated

We compared the capacities of human antisera derived from 40 COVID-19 convalescent donors to block infection by rVSV-SARS-CoV-2 S and authentic SARS-CoV-2 in a microneutralization format. Briefly, pre-titrated amounts of viral particles were incubated with serial dilutions of each antiserum, and target cells were then exposed to the virus-antiserum mixtures. Viral infection was determined by enumerating eGFP-positive cells (rVSV) as above (Fig. 1) or cells immunoreactive with a SARS-CoV-2 nucleocapsid protein-specific antibody (authentic virus) (Fig. 5A). Heatmaps of viral infectivity revealed similar antiserum donor- and dose-dependent neutralization patterns for rVSV-SARS-CoV-2 S and authentic SARS-CoV-2 (Fig. 5B). Comparison of the serum dilutions at half-maximal neutralization derived from logistic curve fits (neutralization IC_50_) revealed a quantitative shift of 0.5–1.0 log_10_ unit towards enhanced neutralization with rVSV-SARS-CoV-2 S (Fig. 5C). The origin of this difference is unclear but may arise from assay-specific differences in the rVSV and authentic virus microneutralization formats employed herein. Nevertheless, the relative potencies of the antisera against rVSV-SARS-CoV-2 S and authentic SARS-CoV-2 were well correlated (R^2^=0.76) (Fig. 5D). In sum, these findings demonstrate the suitability of rVSV-SARS-CoV-2 S for rapid, high-throughout, reporter-based assays of spike glycoprotein-dependent entry and its inhibition.

**Fig 5.**
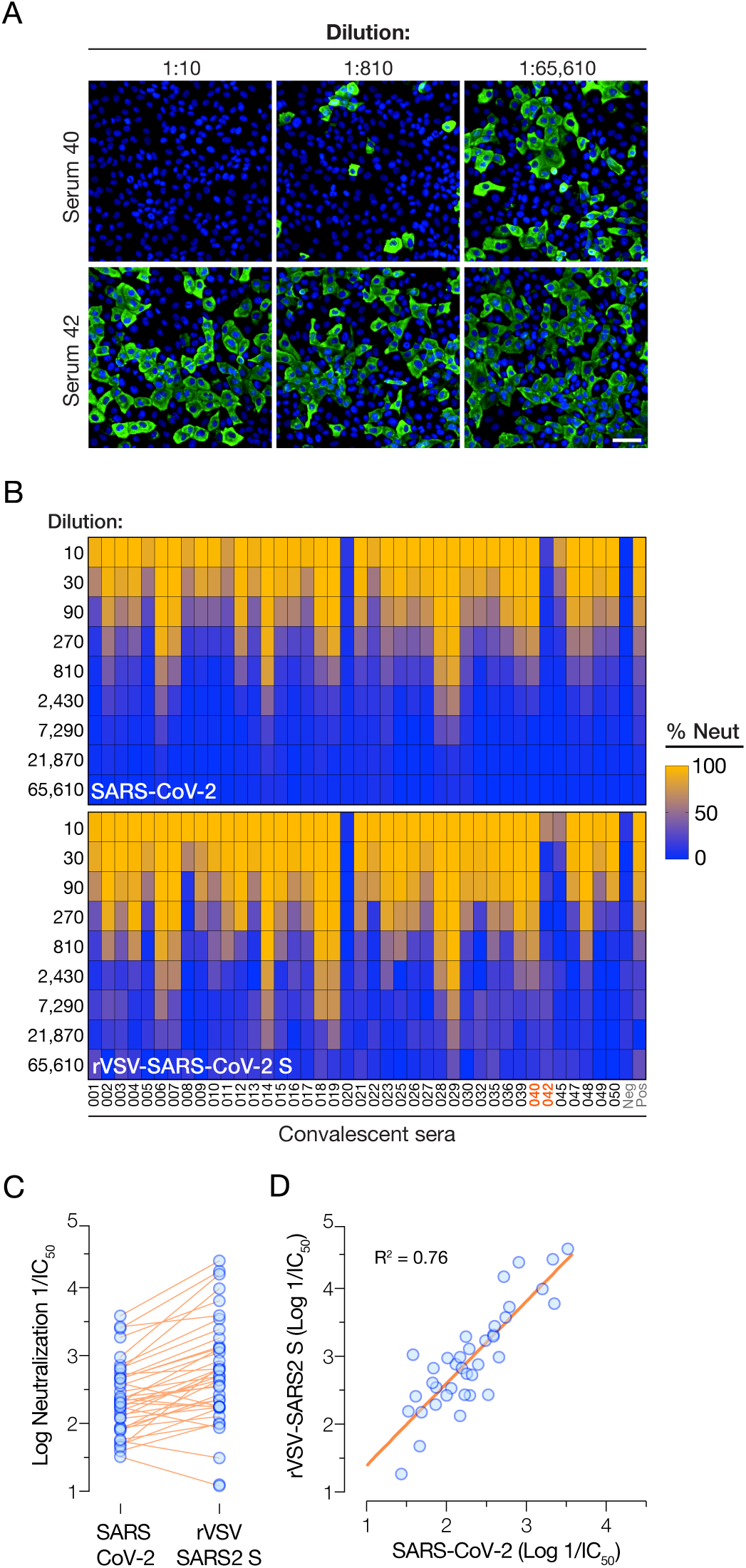
Correlation of convalescent serum-mediated neutralization of rVSV-SARS-CoV-2 S and authentic SARS-CoV-2. **(A)** Pre-titrated amounts of SARS-CoV-2 were incubated with serial 3-fold dilutions of antisera from COVID-19 convalescent donors or negative control at 37°C for 1 h. Virus:serum mixtures were then applied to monolayers of Vero-E6 cells. At 24 h post-infection, cells were fixed, permeabilized and immunostained with a SARS-CoV nucleocapsid-specific antibody. Nuclei were counterstained, infected cells were scored for the presence of nucleocapsid antigen. Representative images from one of the 2 independent experiments are shown (scale bar = 200 µm). **(B)** Pre-titrated amounts of rVSV-SARS-CoV-2 S were incubated with serial 3-fold dilutions of antisera from COVID-19 convalescent patients or negative control at 37°C for 1 h. Virus:serum mixtures was then applied to monolayers of Vero cells. At 16–18 h post-infection, cells were fixed, nuclei were counterstained and infected cells were scored by GFP expression. Heat maps showing % neutralization of authentic SARS-CoV-2 or rVSV-SARS-CoV-2 S by the panel of 40 antisera are shown (Averages of n = 4 from 2 independent experiments). **(C)** Comparison of the neutralizing activities of the antisera (log reciprocal IC50 values) against authentic SARS-CoV-2. and rVSV-SARS-CoV-2 S. **(D)** Linear regression analyses of neutralization IC50 values from panel C.

**Fig S2.**
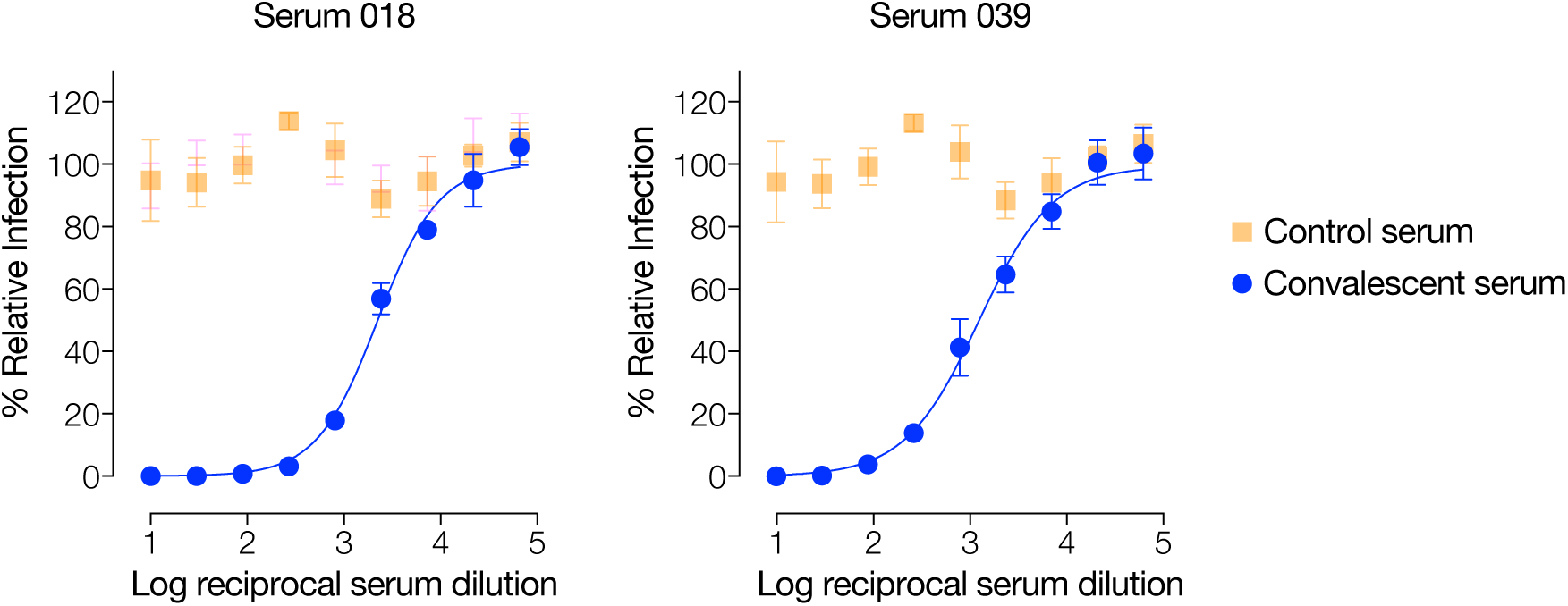
rVSV-SARS-CoV-2 S neutralization dose-curves with antisera from convalescent donors. Pre-titrated amounts of rVSV-SARS-CoV-2 S were incubated with serial 3-fold dilutions of antisera from two COVID-19 convalescent donorsor negative control at 37°C for 1 h. Virus:serum mixtures were then applied to monolayers of Vero cells. At 16-18 h post-infection, cells were fixed, nuclei were counterstained, and infected cells were scored for eGFP expression (Average ± SD, n = 4 from 2 independent experiments).

## Discussion

There is an urgent need for vaccines and therapeutics to prevent and treat COVID-19. The rapid development of SARS-CoV-2 countermeasures is contingent on the availability of robust, scalable, and readily deployable surrogate viral systems to screen antiviral humoral responses and define correlates of immune protection. Such tools would also facilitate the efficient down-selection of candidate antivirals and studies of their mechanisms of action. Here, we describe a highly infectious recombinant vesicular stomatitis virus bearing the SARS-CoV-2 spike glycoprotein S that closely resembles the authentic agent in its entry-related properties. We show that rVSV-SARS-CoV-2 S affords the high-throughput, reporter-based screening of small-molecule and antibody-based inhibitors targeting the viral spike glycoprotein with performance characteristics comparable to those of SARS-CoV-2.

rVSV-SARS-CoV-2 S initially replicated poorly in cell culture following its rescue from plasmids, but we noted accelerated viral growth at passage 5 (Fig. 1). This coincided with the emergence of viral variants bearing S glycoproteins with 21- or 24-amino acid truncations of their cytoplasmic tails. The cytoplasmic tails of the S glycoproteins of SARS-CoV and SARS-CoV-2 are highly similar and carry signals for their retention in the endoplasmic reticulum (ER), including a conserved K×H×× motif located near the C–terminus (McBride et al. 2007; Ujike et al. 2016). 18–19-amino acid deletions in the cytoplasmic tails of SARS-CoV S (Fukushi et al. 2005; Fukushi et al. 2006) and SARS-CoV-2 S (Ou et al. 2020) have been shown to increase the infectivity of single-cycle VSV-S pseudotypes. As previously observed for ER/Golgi-localizing hantavirus glycoproteins (Slough et al. 2019), these deletions likely redistribute S glycoproteins to the cell surface, thereby relieving the spatial mismatch in budding between VSV and SARS-CoV2 (plasma membrane vs. ER, respectively) and enhancing S incorporation into VSV particles.

Accelerated growth by rVSV-SARS-CoV-2 S around passage 5 was accompanied by a marked increase in the occurrence of syncytia (see Fig. 1B) due to S-mediated cell-cell fusion (Fig. S1). This may reflect a functional property of the cytoplasmic tail-deleted S variants, including perturbations in their subcellular localization, as discussed above.

Strikingly, passage 9 stocks and highly infectious viral plaque isolates from these stocks displayed a pattern of spreading infection more typical for rVSVs, with few large syncytia in evidence (Fig. 1B). In this regard, it is tempting to speculate that one or more additional S glycoprotein mutations detected in the passage 5 and 9 viral populations and in the six plaque isolates (Table S1) arose as compensatory changes to suppress the syncytiogenic propensity of the rVSV-encoded SΔ21 glycoprotein spikes. Indeed, several mutations in the S1 NTD and RBD may serve to modulate spike glycoprotein fusogenicity, as also may mutations at the S1– S2 and S2’ cleavage sites (R685G and P812R, respectively) (Table S1). Further, at least one mutation (H655Y), present in five of six rVSV-SARS-CoV-2 S plaque isolates, has arisen during natural SARS-CoV-2 evolution in humans (Yang et al., 2020), during transmission studies in a hamster model (Chan et al., 2020), and possibly during SARS-CoV-2 passage in tissue culture (this report). Our current efforts are aimed at understanding what role(s) these mutations in the S glycoprotein ectodomain play in the maintenance of high levels of rVSV infectivity without the formation of large numbers of syncytia. These findings also highlight a feature of rVSV-SARS-CoV-2 S not shared by any of the viral entry surrogates described to date—its utility for forward genetics. This can be used to dissect structure-function relationships in the SARS-CoV-2 spike glycoprotein and to elucidate the mechanisms of action of spike- or entry-targeted antivirals.

We demonstrate that rVSV-SARS-CoV-2 S can be used to rapidly and faithfully assess the neutralizing activities of large panels of COVID-19 convalescent sera (this report, Fig. 5) and spike-directed mAbs (Walker et al., 2020). We have exploited the fidelity and high throughput of our rVSV-based 384-well plate microneutralization assay to rapidly pre-screen >300 COVID-19 convalescents and identify potential convalescent plasma donors first for the expanded access program and now for an ongoing randomized controlled trial of convalescent plasma therapy (Casadevall and Pirofski, 2020; Clinical Trial - NYU/AECOM, 2020; Chandran & Pirofski, manuscript in preparation). As the COVID-19 pandemic continues apace and the development of plasma-, hyperimmune globulin-, mAb-, and small molecule-based countermeasures accelerates, the need for highly scalable surrogate viral assays continues to mount. The availability of a highly infectious rVSV surrogate that can be scaled up with relative ease to produce standardized batches for antiviral screening and readily deployed in reporter-based microneutralization assays will facilitate these efforts.

## Acknowledgments

We thank I. Gutierrez, E. Valencia, and L. Polanco for laboratory management and technical support. We thank J. McLellan for his generous gifts of wild-type and recombinant stabilized versions of SARS-CoV-2 spike constructs. We also thank F. Krammer, H. Choe and R. Reeves for their generous gifts of SARS-CoV-2 RBD, hACE2 and TMPRSS2 constructs, respectively. This work was supported in part by National Institutes of Health (NIH) grants U19AI142777 and R01AI132633 (to K.C.) and R01AI143453 and R01AI123654 (to L.P.), R01AI125462 (to J.R.L.) and R21AI141367 (to J.P.D). M.E.D. is a Latin American Fellow in the Biomedical Sciences, supported by the Pew Charitable Trusts. C.F. is an NRC Research Associateship Program Fellow. R.H.B.III. and R.J.M. were partially supported by NIH training grant 2T32GM007288-45 (Medical Scientist Training Program) at Albert Einstein College of Medicine.

## Materials and Methods

### Cells

Human hepatoma Huh7.5.1 (received from Dr. Jan Carette; originally from Dr. Frank Chisari) and 293FT (ThermoFisher) cells were cultured in Dulbecco’s Modified Eagle Medium (DMEM high glucose, Gibco) supplemented with 10% heat-inactivated fetal bovine serum (FBS, Atlanta Biologicals), 1% Penicillin/Streptomycin (P/S, Gibco) and 1% Gluta-MAX (Gibco). The African vervet monkey kidney Vero cells and baby hamster kidney BHK21 cells were maintained in DMEM (high glucose) supplemented with 2% heat-inactivated FBS, 1% P/S and 1% Gluta-MAX. Vero-E6 cells were grown in MEM supplemented in 10% FBS and Gentimicine. All cell lines were passaged every 2-3 days using 0.05% Trypsin/EDTA solution (Gibco).

### Convalescent serum samples

Serum samples were collected from healthy adult volunteers residing in Westchester County, NY who had recovered from COVID-19 in April 2020. Patients had reported a positive nasopharyngeal swab by PCR for SARS-CoV-2 during illness and had been asymptomatic for at least 14 days prior to sample collection. After obtaining informed consent, serum was obtained by venipuncture (BD Vacutainer, serum), centrifuged, aliquoted and stored at -80°C prior to use. The sera were heat-inactivated at 56°C for 30 minutes and stored at 4°C prior to analysis. Protocol approval was obtained by the Institutional Review Board (IRB) of the Albert Einstein College of Medicine.

### Generation of rVSV-SARS-CoV-2

A plasmid encoding the VSV antigenome was modified to replace its native glycoprotein, G, with the full-length wild-type *S* glycoprotein gene of the Wuhan-Hu-1 isolate of SARS-CoV-2 (GenBank MN908947.3). The VSV antigenome also encodes for an eGFP reporter gene as a separate transcriptional unit. Plasmid-based rescue of the rVSV was carried out as described previously (Kleinfelter et al., 2015; Whelan et al., 1995; Wong et al., 2010). Briefly, 293FT cells were transfected with the VSV antigenome plasmid along with plasmids expressing T7 polymerase and VSV N, P, M, G and L proteins by using polyethylenimine. Supernatants from the transfected cells were transferred to Huh7.5.1 cells every day (day 2-7 post-transfection) till the appearance of eGFP-positive cells. The poorly spreading virus was initially propagated by cell subculture. RNA was isolated from viral supernatants at different passages and Sanger sequencing was used to verify *S* gene sequences. A passage #9 viral stock was plaque-purified on Vero cells. Supernatants were aliquoted and stored at -80°C.

### Detection of S protein incorporation in rVSV-SARS-CoV-2

High-protein binding 96-well ELISA plates (Corning) were coated with 25 µl of concentrated rVSV-SARS-CoV-2 S or rVSV-EBOV (2.73 µg/ml) overnight at 4°C, and blocked with 3% nonfat dry milk in PBS (PBS-milk) for 1 h at 25°C. Plates were extensively washed and incubated with serum 18, serum 39 or negative serum diluted to 1:100 first then with serial 2-fold dilutions in PBS milk 1% -Tween 0.1% for 1 h at 25°C. Plates were washed three times and incubated with Goat anti-human IgG-HRP (#31410 Invitrogen) diluted 1:3000 (PBS milk 1% -Tween 0.1%) for 1 h at 25°C and detected using 1-Step(tm) Ultra TMB-ELISA Substrate Solution (Thermo Fisher Scientific, Waltham, MA). Plates were read using Cytation5 (BioTek) at 450 nm.

### NH_4_Cl inhibition experiments

Huh7.5.1 cell monolayers were incubated for 1hour with 20– 100 mM NH_4_Cl in DMEM. Next, pre-titrated rVSVs expressing EBOV GP or SARS-CoV-2 S were used to infect cells. Infection was scored 16–18 h later as described above.

### Cathepsin inhibitor experiments

Monolayers of Vero cells pre-treated for 1.5 hours at 37°C with E-64 (37.5 or 75 µM), Z-Phe-tyr-dmk (FYdmk, 18.75 or 37.5 µM), or 1.5% DMSO (vehicle control) were infected with pre-titrated amounts of rVSVs carrying SARS-CoV 2 S, EBOV GP or VSV G. At 1 h post-infection, 20 mM NH_4_Cl was added. Infected cells were fixed 16–18 h later and scored for infection as described above.

### Generation of BHK21-hACE2 and Vero-TMPRSS2 cells

Human ACE2 and TMPRSS2 coding sequences were PCR-amplified from the hACE2 plasmid Addgene #1786 (a generous gift from Hyeryun Choe) and TMPRSS2 plasmid Addgene #53887 (a generous gift from Roger Reeves), respectively and cloned into a retroviral pBabe-puro vector. Retroviruses were produced by transfecting 293FT cells with the hACE2 and TMPRSS2 expressing pBabe-puro plasmids along with those expressing the Moloney murine leukemia virus (MMLV) gag-pol and VSV G proteins. Retroviral supernatants passed through a 0.45-µm filter were used to transduce BHK21 or Vero cells. Transfected cells were selected with puromycin (2 µg/ml).

### Detection of hACE2 in BHK21 transfected cells

To stain for surface-expressed hACE2, BHK21-hACE2 or the control BHK21 cells seeded onto fibronectin-coated glass coverslips were incubated with 0.4 μg/mL of hACE2-specific goat antibody (#AF933, R&D systems) at 4°C in media containing 25 mM HEPES. Next, cells were washed with cold PBS, fixed with 4% paraformaldehyde, and blocked with buffer (2% (w/v) bovine serum albumin, 5% (v/v) glycerol, 0.2% (v/v) Tween20 in Ca^2+^/Mg^2+^-free PBS). Secondary donkey AlexaFluor 594-conjugated anti-goat IgG (#A32758 Invitrogen) was used to detect the hACE2 signal. Coverslips were mounted in Prolong with DAPI (Invitrogen) and imaged on an Axio Observer inverted microscope (Zeiss).

### rVSV-SARS-CoV-2 S microneutralization assay

Serum samples were serially diluted and incubated with virus for 1 h at room temperature. Serum:virus mixtures were then added in duplicate to 384-well plates (Corning) containing Huh7.5.1 cells or 96-well plates (Corning) containing Vero cells. Plates were incubated for 16–18 h at 37°C and 5% CO2. Cells were fixed with 4% paraformaldehyde (Sigma), washed with PBS, and stored in PBS containing Hoechst-33342 (Invitrogen) at a dilution of 1:2,000 in. Viral infectivity was measured by automated enumeration of GFP-positive cells from captured images using a Cytation5 automated fluorescence microscope (BioTek) and analyzed using the Gen5 data analysis software (BioTek). The half-maximal inhibitory concentration (IC_50_) of the mAbs was calculated using a nonlinear regression analysis with GraphPad Prism software.

### SARS-CoV2 RBD expression and purification

The pCAGGS SARS-CoV2 RBD plasmid (a generous gift from Florian Krammer) was used for the expression of recombinant RBD as previously described (Amanat et al., 2020; Stadlbauer et al., 2020). FreeStyle 293F cells (ThermoFisher Scientific) were transfected with the plasmid DNA diluted in PBS (0.67 μg total plasmid DNA per ml of culture) using polyethylenimine (Polysciences, Inc.) at a DNA-to-PEI ratio of 1:3. At 6 days post-transfection, cultures were harvested by centrifugation at 4,000 x g for 20 min, and supernatant was incubated with Ni-NTA resin (GoldBio) for 2 h at 4°C. Resin was collected in columns by gravity flow, washed with a wash buffer (50 mM Tris HCl pH 8.0, 250 mM NaCl, 20 mM Imidazole) and eluted with an elution buffer (50 mM Tris HCl pH 8.0, 250 mM NaCl, 250 mM Imidazole). Eluant was concentrated in Amicon centrifugal units (EMD Millipore) and buffer was exchanged into the storage buffer (50 mM Tris HCl pH 8.0, 250 mM NaCl). Protein was analyzed by SDS-PAGE, aliquoted, and stored at -80°C.

### SARS-CoV2 Spike expression and purification

The pCAGGS SARS-CoV2 plasmid encoding stabilized *S* glycoprotein gene (a generous gift from Jason McLellan) was used for the expression of recombinant S protein as described previously (Wrapp et al., 2020) with several modifications. ExpiCHO-S cells (ThermoFisher) were transiently transfected with plasmid DNA diluted in OptiPRO Serum-Free Medium (0.8 μg total DNA per ml of culture) using ExpiFectamine (ThermoFisher) at a DNA-to-ExpiFectamine ratio of 1:4. At 8 days post-transfection, cultures were harvested by centrifugation at 4,000 x g for 20 min. Clarified supernatant was dialyzed in 50 mM Tris HCl pH 8.0, 250 mM NaCl at a clarified supernatant to dialysis buffer ratio of 1:25 prior affinity chromatography. Dialyzed supernatant was incubated with Ni-NTA resin (GoldBio) for 2 h at 4°C. Resin was collected in columns by gravity flow, washed with wash buffer (50 mM Tris HCl pH 8.0, 250 mM NaCl, 20 mM Imidazole) and eluted with elution buffer (50 mM Tris HCl pH 8.0, 250 mM NaCl, 250 mM Imidazole). Eluant was concentrated in Amicon centrifugal units (EMD Millipore) and exchanged into a storage buffer (50 mM Tris HCl pH 8.0, 250 mM NaCl). Protein was analyzed by SDS-PAGE, aliquoted, and stored at -80°C.

### Anti-hACE2 antibody blocking assay

Huh7.5.1 cells were seeded into a 384-well plate. Nex day, goat anti-human ACE2 antibody (#AF933, R&D Systems) was serially diluted and applied to Huh 7.5.1 cells. After 1 h incubation at 37°C and 5% CO2, cells were infected with rVSV-SARS-CoV-2 S. At 16-18 h post-infection, cells were fixed and scored for infection as described above. Human gamma globulin (009-000-002) purchased from Jackson ImmunoResearch was used as negative control.

### RBD competition assay

Monolayers of Huh7.5.1 cells in a 384-well plate were incubated with serial dilutions of recombinant RBD domain for 1 hour at 37°C and 5% CO2. Cells were then infected with pre-titrated amounts of rVSV-SARS-CoV-2 or rVSV-EBOV GP and scored for infection 16–18 h later.

### S-mediated antibody depletion assay

High-protein binding 96-well ELISA plates (Corning) coated with PBS alone or with 2 µg/ml of SARS-CoV-2 S protein in PBS overnight at 4°C were blocked for 1 hour with 3% nonfat dry milk (Biorad) in PBS. Serum samples diluted in DMEM (1:50 dilution) were serially incubated 4 times for 1 h each at 37°C on S protein-coated or control wells. The depleted sera were tested for their neutralization capacity as described above.

### SARS-CoV-2 stock preparation

Vero-76 cells were inoculated with SARS-CoV-2 (GenBank MT020880.1) at an MOI of 0.01 and incubated at 37°C with 5% CO2 and 80% humidity. At 50 hours post-infection, cells were frozen at -80°C for 1 hour, allowed to thaw at room temperature, and supernatants were collected and clarified by centrifugation at ∼2,500xg for 10 minutes. Clarified supernatant was aliquoted and stored at -80°C. Sequencing data from this virus stock indicated a single mutation in the spike glycoprotein (H655Y) relative to Washington state isolate MT020880.1.

### SARS-CoV-2 neutralization assay

Serially diluted serum samples were mixed with pre-diluted SARS-CoV-2 in infection media (EMEM/2% FBS/Gentimicine) and incubated for 1 hour at 37°C/5% CO2/80% humidity. Virus/serum inoculum was added to Vero-E6 cells, seeded in 96 well plates, at an MOI of 0.4 and incubated for 1 hour at 37°C/5% CO2/80% humidity. Virus/serum inoculum was removed, and cells were washed 1 time with PBS prior to addition of culture media (MEM/10% FBS/Gentimicine). Following 24hr incubation at 37°C/5% CO2/80% humidity, media was removed, and cells were washed 1 time with PBS. PBS was removed and cells were submerged in 10% formalin for 24hrs. Formalin was removed, and cells were washed with PBS prior to permeabilization with 0.2% Triton-X for 10 minutes at room temperature. Cells were blocked for ∼2 h, and then immunostained with SARS-1 nucleocapsid protein-specific antibody (SinoBiologic; 40143-V08B) and Alexa Fluor-488 labeled secondary antibody. Cells were imaged using an Operetta (Perkin Elmer) high-content imaging instrument and infected cells were determined using Harmony Software (Perkin Elmer).

### Syncytia inhibition assay

Vero cells were infected with pre-titrated amounts of rVSV-SARS-CoV-2 S for 2 hrs at 37°C and 5% CO2. Following the removal of virus inocula, cells were washed with PBS to remove any residual virus and indicated dilutions of convalescent sera were applied to the infected cells. Cells were fixed, their nuclei were counterstained, and syncytia formation was imaged by eGFP expression at 16 h post-infection.

### Statistics and reproducibility

All analyses were carried out in GraphPad Prism 8. The n number associated with each dataset in the figures indicates the number of biologically independent samples. The number of independent experiments and the measures of central tendency and dispersion used in each case are indicated in the figure legends.

## Bibliography

Amanat, F., Stadlbauer, D., Strohmeier, S., Nguyen, T.H.O., Chromikova, V., McMahon, M., Jiang, K., Arunkumar, G.A., Jurczyszak, D., Polanco, J., et al. (2020). A serological assay to detect SARS-CoV-2 seroconversion in humans. Nat. Med.

Bosch, B.J., van der Zee, R., de Haan, C.A.M., and Rottier, P.J.M. (2003). The coronavirus spike protein is a class I virus fusion protein: structural and functional characterization of the fusion core complex. J. Virol. 77, 8801–8811.

Caì, Y., Yú, S., Jangra, R.K., Postnikova, E.N., Wada, J., Tesh, R.B., Whelan, S.P.J., Lauck, M., Wiley, M.R., Finch, C.L., et al. (2019). Human, nonhuman primate, and bat cells are broadly susceptible to tibrovirus particle cell entry. Front. Microbiol. 10, 856.

Cao, Y., Su, B., Guo, X., Sun, W., Deng, Y., Bao, L., Zhu, Q., Zhang, X., Zheng, Y., Geng, C., et al. (2020). Potent neutralizing antibodies against SARS-CoV-2 identified by high-throughput single-cell sequencing of convalescent patients’ B cells. Cell.

Casadevall, A., and Pirofski, L. (2020). The convalescent sera option for containing COVID-19. The Journal of Clinical Investigation.

Chan, J.F.-W., Zhang, A.J., Yuan, S., Poon, V.K.-M., Chan, C.C.-S., Lee, A.C.-Y., Chan, W.-M., Fan, Z., Tsoi, H.-W., Wen, L., et al. (2020). Simulation of the clinical and pathological manifestations of Coronavirus Disease 2019 (COVID-19) in golden Syrian hamster model: implications for disease pathogenesis and transmissibility. Clin. Infect. Dis.

Dong, E., Du, H., and Gardner, L. (2020). An interactive web-based dashboard to track COVID-19 in real time. Lancet Infect. Dis. 20, 533–534.

Fukushi, S., Mizutani, T., Saijo, M., Kurane, I., Taguchi, F., Tashiro, M., and Morikawa, S. (2006a). Evaluation of a novel vesicular stomatitis virus pseudotype-based assay for detection of neutralizing antibody responses to SARS-CoV. J. Med. Virol. 78, 1509–1512.

Fukushi, S., Mizutani, T., Saijo, M., Matsuyama, S., Taguchi, F., Kurane, I., and Morikawa, S. (2006b). Pseudotyped vesicular stomatitis virus for functional analysis of SARS coronavirus spike protein. Adv. Exp. Med. Biol. 581, 293–296.

Hoffmann, M., Kleine-Weber, H., Schroeder, S., Krüger, N., Herrler, T., Erichsen, S., Schiergens, T.S., Herrler, G., Wu, N.-H., Nitsche, A., et al. (2020). SARS-CoV-2 Cell Entry Depends on ACE2 and TMPRSS2 and Is Blocked by a Clinically Proven Protease Inhibitor. Cell 181, 271-280.e8.

Jae, L.T., Raaben, M., Riemersma, M., van Beusekom, E., Blomen, V.A., Velds, A., Kerkhoven, R.M., Carette, J.E., Topaloglu, H., Meinecke, P., et al. (2013). Deciphering the glycosylome of dystroglycanopathies using haploid screens for lassa virus entry. Science 340, 479–483.

Jangra, R.K., Herbert, A.S., Li, R., Jae, L.T., Kleinfelter, L.M., Slough, M.M., Barker, S.L., Guardado-Calvo, P., Román-Sosa, G., Dieterle, M.E., et al. (2018). Protocadherin-1 is essential for cell entry by New World hantaviruses. Nature 563, 559–563.

Kapadia, S.U., Rose, J.K., Lamirande, E., Vogel, L., Subbarao, K., and Roberts, A. (2005). Long-term protection from SARS coronavirus infection conferred by a single immunization with an attenuated VSV-based vaccine. Virology 340, 174–182.

Kapadia, S.U., Simon, I.D., and Rose, J.K. (2008). SARS vaccine based on a replication-defective recombinant vesicular stomatitis virus is more potent than one based on a replication-competent vector. Virology 376, 165–172.

Kleinfelter, L.M., Jangra, R.K., Jae, L.T., Herbert, A.S., Mittler, E., Stiles, K.M., Wirchnianski, A.S., Kielian, M., Brummelkamp, T.R., Dye, J.M., et al. (2015). Haploid genetic screen reveals a profound and direct dependence on cholesterol for hantavirus membrane fusion. MBio 6, e00801.

Lei, C., Qian, K., Li, T., Zhang, S., Fu, W., Ding, M., and Hu, S. (2020). Neutralization of SARS-CoV-2 spike pseudotyped virus by recombinant ACE2-Ig. Nat. Commun. 11, 2070.

Letko, M., Marzi, A., and Munster, V. (2020). Functional assessment of cell entry and receptor usage for SARS-CoV-2 and other lineage B betacoronaviruses. Nat. Microbiol. 5, 562–569.

Liu, R., Wang, J., Shao, Y., Wang, X., Zhang, H., Shuai, L., Ge, J., Wen, Z., and Bu, Z. (2018). A recombinant VSV-vectored MERS-CoV vaccine induces neutralizing antibody and T cell responses in rhesus monkeys after single dose immunization. Antiviral Res. 150, 30–38.

Maier, K.E., Jangra, R.K., Shieh, K.R., Cureton, D.K., Xiao, H., Snapp, E.L., Whelan, S.P., Chandran, K., and Levy, M. (2016). A new transferrin receptor aptamer inhibits new world hemorrhagic fever mammarenavirus entry. Mol. Ther. Nucleic Acids 5, e321.

Nie, J., Li, Q., Wu, J., Zhao, C., Hao, H., Liu, H., Zhang, L., Nie, L., Qin, H., Wang, M., et al. (2020). Establishment and validation of a pseudovirus neutralization assay for SARS-CoV-2. Emerg. Microbes Infect. 9, 680–686.

NYU Langone Health / Albert Einstein College of Medicine (2020) Convalescent Plasma to Limit COVID-19 Complications in Hospitalized Patients. ClinicalTrials.gov Identifier: NCT04364737. Retrieved on May, 19th 2020 from https://clinicaltrials.gov/ct2/show/NCT04364737?term=Montefiore&cond=COVID-19&draw=2&rank=2

Ou, X., Liu, Y., Lei, X., Li, P., Mi, D., Ren, L., Guo, L., Guo, R., Chen, T., Hu, J., et al. (2020). Characterization of spike glycoprotein of SARS-CoV-2 on virus entry and its immune cross-reactivity with SARS-CoV. Nat. Commun. 11, 1620.

Pinto, D., Park, Y.-J., Beltramello, M., Walls, A.C., Tortorici, M.A., Bianchi, S., Jaconi, S., Culap, K., Zatta, F., De Marco, A., et al. (2020). Cross-neutralization of SARS-CoV-2 by a human monoclonal SARS-CoV antibody. Nature.

Pu, T., Ding, C., Li, Y., Liu, X., Li, H., Duan, J., Zhang, H., Bi, Y., and Cun, W. (2020). Evaluate severe acute respiratory syndrome coronavirus 2 infectivity by pseudoviral particles. J. Med. Virol.

Raaben, M., Jae, L.T., Herbert, A.S., Kuehne, A.I., Stubbs, S.H., Chou, Y.-Y., Blomen, V.A., Kirchhausen, T., Dye, J.M., Brummelkamp, T.R., et al. (2017). NRP2 and CD63 are host factors for lujo virus cell entry. Cell Host Microbe 22, 688-696.e5.

Rogers, T.F., Zhao, F., Huang, D., Beutler, N., Abbott, R.K., Callaghan, S., Garcia, E., He, W., Hurtado, J., Limbo, O., et al. (2020). Rapid isolation of potent SARS-CoV-2 neutralizing antibodies and protection in a small animal model. BioRxiv.

Shang, J., Ye, G., Shi, K., Wan, Y., Luo, C., Aihara, H., Geng, Q., Auerbach, A., and Li, F. (2020a). Structural basis of receptor recognition by SARS-CoV-2. Nature.

Shang, J., Wan, Y., Luo, C., Ye, G., Geng, Q., Auerbach, A., and Li, F. (2020b). Cell entry mechanisms of SARS-CoV-2. Proc Natl Acad Sci USA.

Stadlbauer, D., Amanat, F., Chromikova, V., Jiang, K., Strohmeier, S., Arunkumar, G.A., Tan, J., Bhavsar, D., Capuano, C., Kirkpatrick, E., et al. (2020). SARS-CoV-2 Seroconversion in Humans: A Detailed Protocol for a Serological Assay, Antigen Production, and Test Setup. Curr. Protoc. Microbiol. 57, e100.

Tan, C.W., Chia, W.N., Chen, M.I.-C., Hu, Z., Young, B.E., Tan, Y.-J., Yi, Y., Lye, D.C., Anderson, D.E., and Wang, L.-F. (2020). A SARS-CoV-2 surrogate virus neutralization test (sVNT) based on antibody-mediated blockage of ACE2-spike (RBD) protein-protein interaction.

Walker, L.M., Wec, A.Z., Wrap, D., Herbert, A.S., Maurer, D.P., Haslwanter, D., Sakharkar, M., Jangra, R.K., Dieterle, M.E., Lilov, A., et al. (2020). Broad sarbecovirus neutralizing antibodies define a key site of vulnerability on the SARS-CoV-2 spike protein. BioRxiv.

Walls, A.C., Park, Y.-J., Tortorici, M.A., Wall, A., McGuire, A.T., and Veesler, D. (2020). Structure, Function, and Antigenicity of the SARS-CoV-2 Spike Glycoprotein. Cell 181, 281-292.e6.

Wang, Q., Zhang, Y., Wu, L., Niu, S., Song, C., Zhang, Z., Lu, G., Qiao, C., Hu, Y., Yuen, K.-Y., et al. (2020). Structural and Functional Basis of SARS-CoV-2 Entry by Using Human ACE2. Cell.

Whelan, S.P., Ball, L.A., Barr, J.N., and Wertz, G.T. (1995). Efficient recovery of infectious vesicular stomatitis virus entirely from cDNA clones. Proc Natl Acad Sci USA 92, 8388–8392.

Wong, A.C., Sandesara, R.G., Mulherkar, N., Whelan, S.P., and Chandran, K. (2010). A forward genetic strategy reveals destabilizing mutations in the Ebolavirus glycoprotein that alter its protease dependence during cell entry. J. Virol. 84, 163–175.

World Health Organization, Situation report-119, Coronavirus disease (COVID-19) Retrieved on May 19th, 2020 from https://www.who.int/docs/default-source/coronaviruse/situation-reports/20200518-covid-19-sitrep-119.pdf?sfvrsn=4bd9de25_4

Wrapp, D., Wang, N., Corbett, K.S., Goldsmith, J.A., Hsieh, C.L., Abiona, O., Graham, B.S., and McLellan, J.S. (2020). Cryo-EM structure of the 2019-nCoV spike in the prefusion conformation. Science 367, 1260–1263.

Wu, F., Zhao, S., Yu, B., Chen, Y.-M., Wang, W., Song, Z.-G., Hu, Y., Tao, Z.-W., Tian, J.-H., Pei, Y.-Y., et al. (2020a). A new coronavirus associated with human respiratory disease in China. Nature 579, 265–269.

Wu, Y., Wang, F., Shen, C., Peng, W., Li, D., Zhao, C., Li, Z., Li, S., Bi, Y., Yang, Y., et al. (2020b). A noncompeting pair of human neutralizing antibodies block COVID-19 virus binding to its receptor ACE2. Science.

Xiong, H.-L., Wu, Y.-T., Cao, J.-L., Yang, R., Ma, J., Qiao, X.-Y., Yao, X.-Y., Zhang, B.-H., Zhang, Y.-L., Hou, W.-H., et al. (2020). Robust neutralization assay based on SARS-CoV-2 S-bearing vesicular stomatitis virus (VSV) pseudovirus and ACE2-overexpressed BHK21 cells. BioRxiv.

Yang, X., Dong, N., Chan, W.-C., and Chen, S. (2020). Identification of super-transmitters of SARS-CoV-2. MedRxiv.

Zhou, P., Yang, X.-L., Wang, X.-G., Hu, B., Zhang, L., Zhang, W., Si, H.-R., Zhu, Y., Li, B., Huang, C.-L., et al. (2020). A pneumonia outbreak associated with a new coronavirus of probable bat origin. Nature 579, 270–273.

Zost, S.J., Gilchuk, P., Chen, R., Case, J.B., Reidy, J.X., Trivette, A., Nargi, R.S., Sutton, R.E., Suryadevara, N., Chen, E.C., et al. (2020). Rapid isolation and profiling of a diverse panel of human monoclonal antibodies targeting the SARS-CoV-2 spike protein. BioRxiv.

